# Evaluation of efficacy of carfentrazone + metsulfuron, sulfosulfuron + metsulfuron and halauxifen against *Rumex* spp. in wheat (*Triticum aestivum*) in Western Haryana

**DOI:** 10.1101/2021.05.02.442319

**Authors:** Sunil Soni, Samunder Singh, Rajbir Garg

## Abstract

*Rumex* spp. is most dominating broad-leaf weed of wheat crop. Complaints of poor efficacy of different herbicides against *Rumex* spp. have reported recently from different locations of Haryana state. Therefore, the present study was carried out under completely randomised design, replicated thrice, using three different herbicides namely carfentrazone + metsulfuron, sulfosulfuron + metsulfuron and halauxifen as treatments applied with three doses against four populations of *Rumex* spp. Plant height, chlorophyll fluorescence, electrical conductivity, mortality percentage and dry weight were recorded as observations. Results indicated that most of the *Rumex* biotypes were found resistant against sulfosulfuron + metsulfuron application. Majority of biotypes were moderately controlled by the application of halauxifen. Carfentrazone + metsulfuron effectively controlled the *Rumex* spp. and provided 70-90 % control to all biotypes at double of the recommended dose. As per results of this study, carfentrazone + metsulfuron can be recommended for control of *Rumex* spp. in wheat.

## INTRODUCTION

Wheat (*Triticum aestivum* L.) is most important food crop of the world. At world level, it is grown in approximately 214.3 million hctare area with production and productivity of 734.1 million tonne and 3425.5 kg/ha, respectively (FAO STAT, 2018). In India, it is second most important food crop after rice cultivated in 30.6 mha with 99.8 mt production and 3220 kg/ha productivity (Anonymous, 2018). Haryna is the major wheat growing state and rice-wheat, cotton-wheat, bajra-wheat are its major crop sequences. Although there are many obstacles in wheat production, but weed infestation is most prominent among them. In severe cases weeds can cause complete failure of crop (Malik and Singh, 1995). Weeds adversely affect the production of crop by competing with it for various growth factors such as water, solar radiation, nutrients *etc*. (Singh *et al*., 2007).

Although there are many methods for weed management, but chemical method is the most popular method that has revolutionized the weed management programme (Yadav and Malik, 2005). Sulfosulfuron is the most important herbicides used in wheat crop mainly in north western regions of India (Chhokar and Sharma, 2008), but it is not effective against several broadleaf weeds like *Rumex dentatus* and *Convolvulus arvensis* (Chhokar *et al*., 2007a). As tank mixed herbicides are found prominent for effective weed management due to different chemistry of component herbicides, so sulfosulfuron is used in combination with metsulfuron for management of complex weed flora of wheat crop. Herbicidal mixtures are helpful in preventing the problem of target site resistance (Paswan *et al.*, 2012).

*Rumex dentatus* is a major weed that adversely affects crop yield (Chhokar *et al.,* 2007b). It is a serious problem of irrigated weed particularly in rice-wheat cropping system in North Western Indo Gangetic plains of India comprising of state of Haryana (Singh *et al*., 1995). This weed has potential to cause yield loss in wheat crop upto an extent of 55 %. Metsulfuron, sulfonylurea herbicide is recommended for its control for a longer period of time. But recently poor efficacy of this herbicide was reported (Chhokar *et al*., 2015; Singh *et al*., 2017). It was observed that carfentrazone + metsulfuron also have effective results against the broadleaf weeds such as *Rumex* spp. and *C. didymus* (Chhokar *et al*., 2015). Halauxifen when applied in combination with florasulum gives satisfactory results against weed flora of wheat crop (Chhokar *et al*., 2015). Keeping in view the above facts and paucity of research on above aspects, the present investigation planned to evaluate the response of carfentrazone + metsulfuron, sulfosulfuron + metsulfuron and halauxifen against *Rumex* spp. in wheat.

## MATERIAL AND METHODS

A pot experiment was carried out in Department of Agronomy, Chaudhary Charan Singh Haryana Agricultural University, under CRD (Completely Randomized Design) with three replications using seeds of four populations of *Rumex* spp. named as HHH (HAU, Hisar), UPH (Ujha, Panipat), JHH (Jind) and JJH (Jhajjar) collected from farmer’s field of mentioned districts. Geographical coordinates of these biotypes are discussed in table 1. Seeds collected from research farm, CCSHAU, Hisar were used as standard check for comparison. Three herbicides carfentrazone + metsulfuron, sulfosulfuron + metsulfuron and halauxifen were taken and applied at three doses (0.5X, X and 2X) to evaluate their efficacy against *Rumex* spp.

**Table 1.** *Rumex* populations collected for pot experiment.

| Populations | Symbol | Longitude | Latitude |
| --- | --- | --- | --- |
| CCS HAU, Hisar | HHH | 75<sup>0</sup>43’’22’ | 29<sup>0</sup>9’’14’ |
| Ujha, Panipat | UPH | 76<sup>0</sup>58’’5’ | 29<sup>0</sup>23’’20’ |
| JInd | JHH | 76<sup>0</sup>18’’54’ | 29<sup>0</sup>18’’56’ |
| Jhajjar | JJH | 76<sup>0</sup>39’’23’ | 28<sup>0</sup>36’’22’ |

Soil samples were taken randomly from five different spots from a depth of 0-15 cm in experimental field before sowing. The samples were dried and shade, ground and sieved through 2 mm sieve and the composite soil samples were subjected to physic-mechanical and chemical analysis to determine the native fertility and texture of soil. Mechanical and chemical analysis of soil is discussed in table 2 and 3, respectively.

**Table 2.** Mechanical analysis of soil.

| Sr. No. | Fraction | Proportion in percentage | Method of determination |
| --- | --- | --- | --- |
|  |  | Hisar sandy Loam |  |
| 1. | Sand | 63.25 | International pipette method (Piper, 1950) |
| 2. | Silt | 25.30 |  |
| 3. | Clay | 11.45 |  |

**Table 3.**
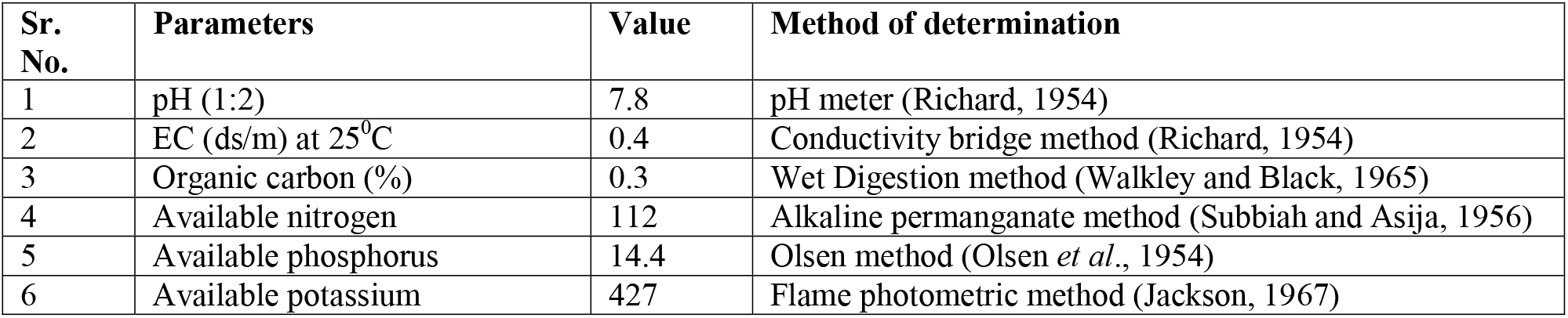
Chemical analysis of soil.

### Pot preparation

Soil from research farm area of agronomy department was used for filling of pots. Soil was finely crushed and air dried so that it can pass through the sieve of 2 mm pore size. Sand and vermin compost are also mixed in soil in such way that soil, sand and vermin-compost are in ratio of 2:3:1 and pots were filled using this mixture. A regular application of water was given to pots for maintaining their mixture level. A depth of 3-4 cm was maintained for sowing and spraying was done by Knapsack sprayer having flat fan nozzle delivering 300l/ha spray volume at 40 psi pressure. Untreated check was maintained for comparison for comparison of results.

### Observations recorded on *Rumex* spp

#### Plant height

A scale was used for measuring the plant height with greater accuracy. The plant height (cm) was recorded from top of soil surface to the tip of fully opened leaf at 4 week after treatment (WAT).

#### Electrical conductivity

The value of EC was taken with the help of EC meter at 4 WAT. For this, beakers containing weed samples were subjected to boiling at specific temperature for dissolution of salts of weed samples in distilled water. Then the EC of this water is recorded.

#### Chlorophyll Fluorescence (Fv/Fm)

This reading was taken with the help of chlorophyll meter at 7 days after treatment (DAT).

#### Per cent control

Mortality percentage was recorded at 4 WAT in scale of 0-100 (0 means no control and 100 means complete control of *Rumex* spp.). The observations were recorded by comparing each treatment with control.

#### Dry weight of weeds

The weed samples were harvested and dried firstly and then final drying was done inside oven at 65°C till a constant weight was achieved. Then the weight of these dried samples were taken and expressed as g/m^2^.

#### Statistical analysis

Analyses of all observations were done with the help of OPSTAT software. Arcsine transformation was also used for analysis of observations of mortality percentage.

## RESULT

### Czarfentrazone + metsulfuron dose response study

Table 4. presents the data on various parameters of *Rumex* biotypes as affected by application of carfentrazone + metsulfuron such as plant height, electrical conductivity and per cent control at 4 WAT, chlorophyll fluorescence at 7 DAT and dry weight at harvesting. When data was averaged over carfentrazone + metsulfuron doses, significantly higher plant height (23.6 cm) and higher chlorophyll fluorescence (0.70 fv/fm) was observed in UPH followed by JHH and other biotypes. Half dose of carfentrazone + metsulfuron resulted in 10.6 % higher plant height and 21.8 % higher plant chlorophyll fluorescence over recommended dose whereas double dose resulted in 5.8 % lower plant height and 10.9 % lower plant chlorophyll fluorescence than recommended dose, when data was averaged over all biotypes. Significantly lower EC (ds/m) was observed in UPH (0.32) which was statistically similar with JJH (0.33) and JHH (0.34) followed by HHH (0.37) at 4 WAT

**Table 4.** Plant height, chlorophyll fluorescence, electrical conductivity, mortality percentage and dry weight of *Rumex* biotypes as influenced by Carfentrazone + metsulfuron application

|  | Carfentrazone + metsulfuron (g/ha) |  |  |  |  |  |  |  |  |  |  |  |  |  |  |  |  |  |  |  |  |  |  |  |  |
| --- | --- | --- | --- | --- | --- | --- | --- | --- | --- | --- | --- | --- | --- | --- | --- | --- | --- | --- | --- | --- | --- | --- | --- | --- | --- |
| Populations | Plant height (cm) at 4 WAT |  |  |  |  | Chlorophyll fluorescence (Fv/Fm) at 7 DAT |  |  |  |  | Electrical conductivity (Ds/m) at 4 WAT |  |  |  |  | Mortality percentage At 4 WAT |  |  |  |  | Dry weight (g/pot) at harvesting |  |  |  |  |
|  | 0 | 12.5 | 25 | 50 | Mean | 0 | 12.5 | 25 | 50 | Mean | 0 | 12.5 | 25 | 50 | Mean | 0 | 12.5 | 25 | 50 | Mean | 0 | 12.5 | 25 | 50 | Mean |
| HHH | 26.3 | 19.0 | 18.3 | 17.3 | 20.3 | 0.85 | 0.63 | 0.50 | 0.43 | 0.60 | 0.02 | 0.40 | 0.51 | 0.52 | 0.37 | 0 (0) | 67 (85) | 76 (92) | 85 (98) | 57 (69) | 2.47 | 0.47 | 0.27 | 0.13 | 0.83 |
| UPH | 27.3 | 24.3 | 22.3 | 20.3 | 23.6 | 0.91 | 0.71 | 0.61 | 0.56 | 0.70 | 0.03 | 0.32 | 0.44 | 0.47 | 0.32 | 0 (0) | 49 (57) | 63 (78) | 82 (95) | 48 (58) | 3.23 | 1.03 | 0.67 | 0.20 | 1.28 |
| JHH | 27.7 | 22.0 | 20.3 | 19.3 | 22.3 | 0.91 | 0.69 | 0.52 | 0.49 | 0.65 | 0.03 | 0.33 | 0.48 | 0.53 | 0.34 | 0 (0) | 51 (60) | 72 (87) | 81 (97) | 51 (61) | 2.83 | 1.13 | 0.40 | 0.10 | 1.12 |
| JJH | 21.6 | 18.3 | 14.7 | 14.3 | 17.3 | 0.84 | 0.63 | 0.55 | 0.51 | 0.63 | 0.03 | 0.35 | 0.44 | 0.49 | 0.33 | 0 (0) | 76 (91) | 89 (100) | 89 (100) | 64 (73) | 2.40 | 0.47 | 0.20 | 0.13 | 0.80 |
| Mean | 25.8 | 20.9 | 18.9 | 17.8 |  | 0.88 | 0.67 | 0.55 | 0.49 |  | 0.03 | 0.35 | 0.47 | 0.50 |  | 0 (0) | 61 (73) | 75 (89) | 85 (98) |  | 2.73 | 0.78 | 0.38 | 0.14 |  |
|  | CD (P=0.05) |  |  |  | SEm | CD (P=0.05) |  |  |  | SEm | CD (P=0.05) |  |  |  | SEm | CD (P=0.05) |  |  |  | SEm | CD (P=0.05) |  |  |  | SEm |
| Population | 0.9 |  |  |  | 0.31 | 0.02 |  |  |  | 0.008 | 0.02 |  |  |  | 0.007 | 6 |  |  |  | 2.1 | 0.13 |  |  |  | 0.04 |
| Herbicide | 0.9 |  |  |  | 0.31 | 0.02 |  |  |  | 0.008 | 0.02 |  |  |  | 0.007 | 6 |  |  |  | 2.1 | 0.13 |  |  |  | 0.04 |
| Population x herbicide | NS |  |  |  | 0.62 | 0.05 |  |  |  | 0.016 | 0.04 |  |  |  | 0.014 | 12 |  |  |  | 4.2 | 0.25 |  |  |  | 0.09 |

Original figures in parenthesis were subjected to angular transformation. WAT, weeks after treatment; DAT, days after treatment

Similarly lower mortality percentage was recorded in UPH (48) followed by JJH (51), HHH (57) and JJH (64) at 4 WAT (during mean data over herbicide doses). Half dose of carfentrazone + metsulfuron resulted in 18.7 % lower mortality over recommended dose, whereas double dose resulted in 13.3 % higher mortality than recommended dose at 4 WAT. When data was averaged over carfentrazone + metsulfuron doses, significantly higher dry weight (g/pot) was recorded in UPH (1.28), followed by JHH (1.12), HHH (0.83) and JJH (0.80) at harvesting (120 DAS). Half dose of carfentrazone + metsulfuron resulted in 105.0 % higher dry weight over recommended dose, whereas double dose resulted in 63.2 % lower dry weight than recommended dose at harvesting over all biotypes.

### Sulfosulfuron + metsulfuron dose response study

Table 5. presents the data on plant height, chlorophyll fluorescence, electrical conductivity, per cent control and dry weight of *Rumex* biotypes as affected by the application of sulfosulfuron + metsulfuron, respectively at 4 WAT, 7 DAT, 4 WAT, 4 WAT and harvesting. When data was averaged over sulfosulfuron + metsulfuron doses, significantly higher plant height (cm) was observed in UPH (25.0) followed by JHH (23.9), HHH (20.1) and JJH (17.6). Similarly higher chlorophyll fluorescence (Fv/Fm) was observed in UPH and JHH (0.82) followed by JJH and HHH (0.70). Half dose of sulfosulfuron + metsulfuron resulted in 11.5 % higher plant height and 2.7 % higher plant chlorophyll fluorescence over recommended dose, whereas double dose resulted in 7.0% lower plant height and 10.8 % lower chlorophyll fluorescence than recommended dose, when data was averaged over all biotypes. Significantly lower EC (ds/m) was observed in UPH and JHH (0.07) followed by JJH and HHH (0.24) after boiling at 4 WAT (mean data over herbicide doses).

**Table 5.**
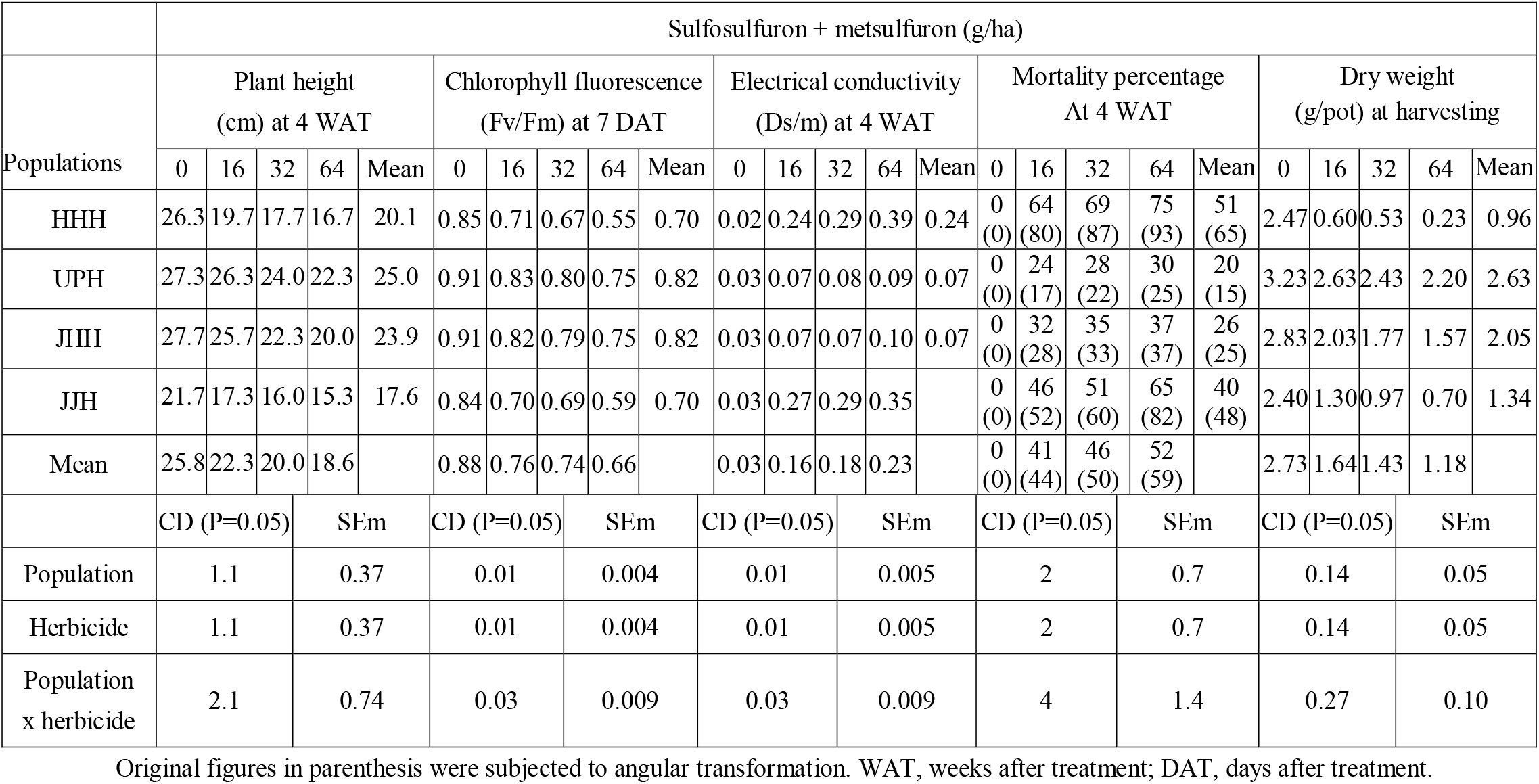
Plant height, chlorophyll fluorescence, electrical conductivity, mortality percentage and dry weight of *Rumex* biotypes as influenced by Sulfosulfuron + metsulfuron application.

|  | Sulfosulfuron + metsulfuron (g/ha) |  |  |  |  |  |  |  |  |  |  |  |  |  |  |  |  |  |  |  |  |  |  |  |  |
| --- | --- | --- | --- | --- | --- | --- | --- | --- | --- | --- | --- | --- | --- | --- | --- | --- | --- | --- | --- | --- | --- | --- | --- | --- | --- |
|  | Plant height<br>(cm) at 4 WAT |  |  |  |  | Chlorophyll fluorescence<br>(Fv/Fm) at 7 DAT |  |  |  |  | Electrical conductivity<br>(Ds/m) at 4 WAT |  |  |  |  | Mortality percentage<br>At 4 WAT |  |  |  |  | Dry weight<br>(g/pot) at harvesting |  |  |  |  |
| Populations | 0 | 16 | 32 | 64 | Mean | 0 | 16 | 32 | 64 | Mean | 0 | 16 | 32 | 64 | Mean | 0 | 16 | 32 | 64 | Mean | 0 | 16 | 32 | 64 | Mean |
| HHH | 26.3 | 19.7 | 17.7 | 16.7 | 20.1 | 0.85 | 0.71 | 0.67 | 0.55 | 0.70 | 0.02 | 0.24 | 0.29 | 0.39 | 0.24 | 0<br>(0) | 64<br>(80) | 69<br>(87) | 75<br>(93) | 51<br>(65) | 2.47 | 0.60 | 0.53 | 0.23 | 0.96 |
| UPH | 27.3 | 26.3 | 24.0 | 22.3 | 25.0 | 0.91 | 0.83 | 0.80 | 0.75 | 0.82 | 0.03 | 0.07 | 0.08 | 0.09 | 0.07 | 0<br>(0) | 24<br>(17) | 28<br>(22) | 30<br>(25) | 20<br>(15) | 3.23 | 2.63 | 2.43 | 2.20 | 2.63 |
| JHH | 27.7 | 25.7 | 22.3 | 20.0 | 23.9 | 0.91 | 0.82 | 0.79 | 0.75 | 0.82 | 0.03 | 0.07 | 0.07 | 0.10 | 0.07 | 0<br>(0) | 32<br>(28) | 35<br>(33) | 37<br>(37) | 26<br>(25) | 2.83 | 2.03 | 1.77 | 1.57 | 2.05 |
| JJH | 21.7 | 17.3 | 16.0 | 15.3 | 17.6 | 0.84 | 0.70 | 0.69 | 0.59 | 0.70 | 0.03 | 0.27 | 0.29 | 0.35 |  | 0<br>(0) | 46<br>(52) | 51<br>(60) | 65<br>(82) | 40<br>(48) | 2.40 | 1.30 | 0.97 | 0.70 | 1.34 |
| Mean | 25.8 | 22.3 | 20.0 | 18.6 |  | 0.88 | 0.76 | 0.74 | 0.66 |  | 0.03 | 0.16 | 0.18 | 0.23 |  | 0<br>(0) | 41<br>(44) | 46<br>(50) | 52<br>(59) |  | 2.73 | 1.64 | 1.43 | 1.18 |  |
|  | CD (P=0.05) |  |  | SEm |  | CD (P=0.05) |  |  | SEm |  | CD (P=0.05) |  |  | SEm |  | CD (P=0.05) |  |  | SEm |  | CD (P=0.05) |  |  | SEm |  |
| Population | 1.1 |  |  | 0.37 |  | 0.01 |  |  | 0.004 |  | 0.01 |  |  | 0.005 |  | 2 |  |  | 0.7 |  | 0.14 |  |  | 0.05 |  |
| Herbicide | 1.1 |  |  | 0.37 |  | 0.01 |  |  | 0.004 |  | 0.01 |  |  | 0.005 |  | 2 |  |  | 0.7 |  | 0.14 |  |  | 0.05 |  |
| Population<br>x herbicide | 2.1 |  |  | 0.74 |  | 0.03 |  |  | 0.009 |  | 0.03 |  |  | 0.009 |  | 4 |  |  | 1.4 |  | 0.27 |  |  | 0.10 |  |

Original figures in parenthesis were subjected to angular transformation. WAT, weeks after treatment; DAT, days after treatment.

Significantly lower mortality (%) was recorded in UPH (20) followed by JHH (26), JJH (40) and HHH (51). Half dose of sulfosulfuron + metsulfuron resulted in 10.9% lower mortality over recommended dose, whereas double dose resulted in 13% higher mortality than recommended dose at 4 WAT. When data was averaged over sulfosulfuron + metsulfuron doses, significantly higher dry weight (g/pot) was recorded in UPH (2.63) followed by JHH (2.05), JJH (1.34) and HHH (0.96) at harvesting (120 DAS). Half dose of sulfosulfuron + metsulfuron resulted in 14.7% higher dry weight over recommended dose, whereas double dose resulted in 17.5% lower dry weight than recommended dose at harvesting (120 DAS), over all biotypes.

### Halauxifen dose-response studies

Table 6 presents the data on various parameter of *Rumex* biotypes as affected by the application of halauxifen such as plant height, electrical conductivity and per cent control at 4 (WAT), chlorophyll fluorescence at 7 (DAT) and dry weight at harvesting. When data was averaged over halauxifen doses, significantly higher plant height (cm) was recorded in UPH (23.7) followed by JHH (22.7), HHH (21.0) and JJH (17.3) at 4 WAT. Similarly higher plant chlorophyll fluorescence (Fv/Fm) was observed in UPH (0.74) which was statistically similar with JJH (0.72) but significantly higher than other biotypes at 7 DAT. Half dose of halauxifen resulted in 8.1% higher plant height and 6.3% higher plant chlorophyll fluorescence over recommended dose, whereas double dose resulted in 9.1% lower plant height and 15.9% lower plant chlorophyll fluorescence than recommended dose, when data was averaged over all biotypes. Significantly lower EC (ds/m) was observed in JJH (0.20) which was statistically similar with UPH (0.21) but significantly lower than other biotypes at 4 WAT (mean data over herbicide doses).

**Table 6.** Plant height, chlorophyll fluorescence, electrical conductivity, mortality percentage and dry weight of

|  | Halauxifen (g/ha) |  |  |  |  |  |  |  |  |  |  |  |  |  |  |  |  |  |  |  |  |  |  |  |  |
| --- | --- | --- | --- | --- | --- | --- | --- | --- | --- | --- | --- | --- | --- | --- | --- | --- | --- | --- | --- | --- | --- | --- | --- | --- | --- |
| Populations | Plant height<br>(cm) at 4 WAT |  |  |  |  | Chlorophyll fluorescence<br>(Fv/Fm) at 7 DAT |  |  |  |  | Electrical conductivity<br>(Ds/m) at 4 WAT |  |  |  |  | Mortality percentage<br>At 4 WAT |  |  |  |  | Dry weight<br>(g/pot) at harvesting |  |  |  |  |
|  | 0 | 6 | 12 | 24 | Mean | 0 | 6 | 12 | 24 | Mean | 0 | 6 | 12 | 24 | Mean | 0 | 6 | 12 | 24 | Mean | 0 | 6 | 12 | 24 | Mean |
| HHH | 26.3 | 21.0 | 19.3 | 17.3 | 21.0 | 0.85 | 0.58 | 0.53 | 0.39 | 0.59 | 0.02 | 0.33 | 0.34 | 0.43 | 0.28 | 0<br>(0) | 54<br>(65) | 56<br>(68) | 65<br>(82) | 44<br>(54) | 2.47 | 0.90 | 0.77 | 0.60 | 1.18 |
| UPH | 27.3 | 24.7 | 22.3 | 20.3 | 23.7 | 0.91 | 0.72 | 0.69 | 0.65 | 0.74 | 0.03 | 0.26 | 0.27 | 0.30 | 0.21 | 0<br>(0) | 49<br>(57) | 52<br>(62) | 56<br>(68) | 39<br>(47) | 3.23 | 1.50 | 1.33 | 0.97 | 1.76 |
| JHH | 27.7 | 23.0 | 21.0 | 19.0 | 22.7 | 0.91 | 0.68 | 0.61 | 0.43 | 0.66 | 0.03 | 0.27 | 0.34 | 0.41 | 0.26 | 0<br>(0) | 53<br>(63) | 55<br>(67) | 59<br>(73) | 42<br>(51) | 2.83 | 1.00 | 0.93 | 0.83 | 1.40 |
| JJH | 21.7 | 16.3 | 16.0 | 15.0 | 17.3 | 0.84 | 0.71 | 0.68 | 0.67 | 0.72 | 0.03 | 0.21 | 0.25 | 0.30 | 0.20 | 0<br>(0) | 43<br>(47) | 52<br>(62) | 53<br>(63) | 37<br>(43) | 2.40 | 1.30 | 0.93 | 0.83 | 1.37 |
| Mean | 25.8 | 21.3 | 19.7 | 17.9 |  | 0.88 | 0.67 | 0.63 | 0.53 |  | 0.03 | 0.27 | 0.30 | 0.36 |  | 0<br>(0) | 50<br>(58) | 54<br>(65) | 58<br>(72) |  | 2.73 | 1.18 | 0.99 | 0.81 |  |
|  | CD (P=0.05) |  |  | SEm |  | CD (P=0.05) |  |  | SEm |  | CD (P=0.05) |  |  | SEm |  | CD (P=0.05) |  |  | SEm |  | CD (P=0.05) |  |  | SEm |  |
| Population | 0.8 |  |  | 0.26 |  | 0.02 |  |  | 0.007 |  | 0.01 |  |  | 0.004 |  | 3 |  |  | 1.0 |  | 0.13 |  |  | 0.05 |  |
| Herbicide | 0.8 |  |  | 0.26 |  | 0.02 |  |  | 0.007 |  | 0.01 |  |  | 0.004 |  | 3 |  |  | 1.0 |  | 0.13 |  |  | 0.05 |  |
| Population<br>x herbicide | NS |  |  | 0.53 |  | 0.04 |  |  | 0.014 |  | 0.02 |  |  | 0.007 |  | NS |  |  | 2.0 |  | 0.26 |  |  | 0.09 |  |

Original figures in parenthesis were subjected to angular transformation. WAT, weeks after treatment; DAT, days after treatment.

Significantly lower mortality (%) was recorded in JJH (37) followed by UPH (39), JHH (42) and HHH (44) at 4 WAT (mean data over herbicide doses). Half dose of halauxifen resulted in 7.4% lower mortality over recommended dose, whereas double dose resulted in 7.4% higher mortality than recommended dose at 4 WAT. When data was averaged over halauxifen doses, significantly higher dry weight (g/pot) was recorded in UPH (1.76) followed by JHH (1.40), JJH (1.37) and HHH (1.18) at harvesting (120 DAS). Half dose of halauxifen resulted in 19.2% higher dry weight over recommended dose, whereas double dose resulted in 18.2% lower dry weight than recommended dose at harvesting (120 DAS), over all biotypes.

## DISSUSSION

### Carfentrazone + metsulfuron dose response study

All of the tested biotypes were found sensitive to the application of carfentrazone + metsulfuron at 4 WAT. At the double of the recommended dose of carfentrazone + metsulfuron, 80-90 % visual control was observed in all biotypes. These results collaborate with the finding of Chhokar *et al*. (2017), Singh *et al*. (2017) and Paswan *et al*. (2012). Carfentrazone + metsulfuron application resulted in lower value of chlorophyll fluorescence mainly due to inhibition of photosystem II. Vershney *et al.* (2012) also observed a reduction in chlorophyll fluorescence value at 1 and 2 DAT in herbicide treated plants.

### Sulfosulfuron + metsulfuron dose response study

Majority of biotypes are highly tolerant against application of sulfosulfuron + metsulfuron at half of the recommended dose, recommended dose and double of the recommended dose. Sulfosulfuron + metsulfuron provided only 30% and 37% mortality in UPH and JHH, respectively at double of the recommended dose at 4 WAT. Singh *et al.* (2017) also reported the low efficacy of sulfosulfuron + metsulfuron against *Rumex* spp. and other broadleaf weeds. These results vary with finding of Bhullar *et al.* (2012) and it may be due to the development of resistance in referred biotypes. UPH biotype attained higher plant height, chlorophyll fluorescence, dry weight and lowest EC, thus confirming the poor efficacy of herbicide.

### Halauxifen dose response study

*Rumex* biotypes were moderately tolerant to halauxifen at recommended dose. It provided 50-70% mortality in all biotypes at recommended dose. Improvement in efficacy was observed at double of the recommended dose of herbicide but still below the level required for its field applicability. Lower control could be due to the lower availability of lethal dose of herbicide to cause satisfactory control. Halauxifen gives effective control of *Rumex* spp. or other broad leaf weeds when it is applied in combination with florasulam (Chhokar *et al*., 2015). The reduction in Fv/Fm value by the application of halauxifen is in the harmony with the findings of Menegat *et al.* (2011).

## CONCLUSION

This experiment has shown the efficacy of carfentrazone + metsulfuron, sulfosulfuron + metsulfuron and halauxifen on four biotypes of *Rumex* spp. in wheat. Among all biotypes, highest emergence percentage was obtained in UPH biotype which implies that good control as to arrest the seed formation would reduce the carry over weed infestation in the next season and could be used as a tool in resistance management in this weed. Sulfosulfuron + metsulfuron provided negligible control to all biotypes which shows that majority of biotypes have attained resistance against sulfosulfuron + metsulfuron. Low efficacy of sulfosulfuron + metsulfuron also reflected in terms of higher value of chlorophyll fluorescence, plant height and dry weight and lower value of electrical conductivity. Carfentrazone + metsulfuron provided 80-90% visual control to al biotypes which shows its good efficacy against Rumex spp. Moderate efficacy of halauxifen was observed at recommended dose which is further improved at double of the recommended dose of herbicide but still below the level required for its field applicability. After comparing the results of carfentrazone + metsulfuron, sulfosulfuron + metsulfuron and halauxifen, it is concluded that carfentrazone + metsulfuron have potential to provide satisfactory control to all biotypes of *Rumex* spp., therefore it can be recommended to farmers for effective control of *Rumex* spp.

## Acknowledgements

Funding and competing interests are not found in this study.

